# Fluidic shear stress alters clathrin dynamics and vesicle formation in endothelial cells

**DOI:** 10.1101/2024.01.02.572628

**Authors:** Tomasz J. Nawara, Elizabeth Sztul, Alexa L. Mattheyses

## Abstract

Endothelial cells (ECs) experience a variety of highly dynamic mechanical stresses. Among others, cyclic stretch and increased plasma membrane tension inhibit clathrin-mediated endocytosis (CME) in non-ECs cells. How ECs overcome such unfavorable, from biophysical perspective, conditions and maintain CME remains elusive. Previously, we have used simultaneous two-wavelength axial ratiometry (STAR) microscopy to show that endocytic dynamics are similar between statically cultured human umbilical vein endothelial cells (HUVECs) and fibroblast-like Cos-7 cells. Here we asked whether biophysical stresses generated by blood flow could favor one mechanism of clathrin-coated vesicle formation to overcome environment present in vasculature. We used our data processing platform – DrSTAR – to examine if clathrin dynamics are altered in HUVECs grown under fluidic sheer stress (FSS). Surprisingly, we found that FSS led to an increase in clathrin dynamics. In HUVECs grown under FSS we observed a 2.3-fold increase in clathrin-coated vesicle formation and a 1.9-fold increase in non-productive flat clathrin lattices compared to cells grown in static conditions. The curvature-positive events had significantly delayed curvature initiation in flow-stimulated cells, highlighting a shift toward flat-to-curved clathrin transitions in vesicle formation. Overall, our findings indicate that clathrin dynamics and CCV formation can be modulated by the local physiological environment and represents an important regulatory mechanism.

**Significance:** Targeted nanomedicine holds a great promise of improved drug bioavailability and specificity. While some cargoes must cross the blood-tissue barrier, the understanding of endocytic pathways in the context of vasculature is limited, which is an obstacle to targeted nanomedicine delivery. In this pilot study we show that the physiological local vascular environment must be considered in the context of internalization of growth factors, membrane proteins, therapeutics, or pathogens. Studies in non-ECs or ECs not cultured under fluidic shear stress do not properly recapitulate clathrin dynamics and will lead to incorrect conclusions.

## Introduction

Endothelial cells line the lumen of blood vessels where they locally experience highly dynamic mechanical stresses, differing in magnitude and type between arteries, veins and capillaries (Trimm and Red-Horse, 2023). These can be divided into contact-derived stresses, originating from vessel topography, curvature, and stiffness, and flow-derived stresses such as fluidic shear stress (FSS), pressure, and tensile strain caused by the blood flow (Dessalles et al., 2021). It has been shown in non-endothelial cells that intrinsic plasma membrane curvature dramatically impacts the proteome required for endocytosis (Cail et al., 2022), while stiffer substrates (Baschieri et al., 2018), cell squeezing, and increased osmotic pressure inhibit endocytosis (Ferguson et al., 2017). The mechanical impact on endocytosis has been recently reviewed (Joseph and Liu, 2020). Additionally, FSS has been show to slow down the internalization rates of polymer nanocarriers targeted to ICAM-1 in endothelial cells (Bhowmick et al., 2012). From a biophysical perspective, the environment present in vasculature seems unfavorable, and capable of abolishing clathrin-mediated endocytosis (CME). Yet, CME is an essential cellular process for endothelial cells that facilitates the internalization of many cargoes, such as vascular endothelial growth factor and vascular endothelial cadherin, and can be hijacked by pathogens such as COVID-19 (Herrscher et al., 2020; Inoue et al., 2007; Katsuno-Kambe and Yap, 2020; Nanes et al., 2012). Thus, CME is indispensable for vascular development and homeostasis (Gaengel and Betsholtz, 2013; Genet et al., 2019; Jones and Shusta, 2007; Potente et al., 2011). It has not yet been determined how endothelial cells preserve CME in their physiological environment and during pathophysiological conditions. Overall, the assumption that CME is a universal and homogenous process is outdated (Chen and Schmid, 2020; Nawara et al., 2022), and it has been highlighted that distinct clathrin-coated structures differentially regulate signaling outcome. Though often studied in “model” fibroblast cell lines, CME must be studied in a cell type and context-dependent manner to reveal context specific regulatory dynamics (Sigismund et al., 2021). Overall, our understanding of endocytic pathways in the context of vasculature is limited, which presents an obstacle to targeted nanomedicine delivery (Rennick et al., 2021).

CME is mediated by the trimeric coat protein clathrin, which polymerizes at the plasma membrane to form vesicles and flat lattices (Kaksonen and Roux, 2018). Depending on the cell line or tissue, clathrin generates clathrin-coated structures of various shapes and sizes, with only a fraction contributing to cargo uptake (Jones and Shusta, 2007; Sochacki and Taraska, 2019). Previously, we used a total internal reflection fluorescence (TIRF)-based technique called simultaneous two-wavelength axial ratiometry (STAR) microscopy to define how clathrin-coated vesicles form in human umbilical vein endothelial cells (HUVECs) grown under static conditions (Nawara et al., 2022). STAR has the power to capture the dynamics and axial (*z*) distribution of clathrin in living cells with a nanoscale axial resolution (Nawara et al., 2023; Nawara and Mattheyses, 2023). Our previous research showed that clathrin dynamics are similar between HUVECs cultured under static conditions and non-endothelial green monkey kidney fibroblast-like (Cos-7) cells. In both cell types we found multiple productive routes of CME mediated by three modes of vesicle formation: nucleation – where endocytosis begins independent of clathrin; constant curvature – where clathrin accumulation and vesicle formation are coinciding; and flat-to-curved transition – where pre-assemblies of flat clathrin lattices transition into clathrin-coated vesicles (Nawara et al., 2022).

Here, we investigated how endothelial cells preserve endocytic dynamics in a biophysically unfavorable environment. HUVECs were exposed to FSS, mimicking a naturally occurring mechanical stress caused by the blood flow, or cultured statically as a control. We then quantified the number and proportion of productive and nonproductive plasma membrane associated clathrin accumulations. For productive events, we compared clathrin-coated vesicle formation dynamics and classified the mode. We show that HUVECs exposed to FSS had a 2.3-fold increase in clathrin-coated vesicle formation and a 1.9-fold increase in non-productive flat clathrin lattices compared to cells grown in static conditions. Moreover, in the long-lasting events (>50 s) we observed a switch in mode of clathrin-coated vesicle formation, with a preference for flat clathrin platform preassembly before vesicle initiation. This data shows that FSS serves as a mechanical stimulus that impacts CME, thereby preserving endocytosis by offsetting the inhibitory effects of other mechanical stresses.

## Materials and methods

### Automated data processing

Raw STAR microscopy data was processed and Δz was calculated using a high-throughput custom-written MATLAB software – DrSTAR (Nawara et al., 2023). Detection, tracking and classification of endocytic events was achieved using CMEanalysis (Aguet et al., 2013), as previously described (Nawara et al., 2022).

### Cell culture

Pooled human umbilical vein endothelial cells (Lonza, C2519A or Sigma, 200P-05N) were cultured at 37 °C and 5% CO_2_ and per manufacture recommendations. EBM-2 media (Lonza, CC-3156) supplemented with EGM-2 SingleQuots (Lonza, CC-4176) or Vascular Cell Basal Medium (ATCC, PCS-100-030) supplemented with Endothelial Cell Growth Kit-VEGF (ATCC, PCS-100-041) were used. Cells between passages 3-6 were used for the experiments.

### Fluidic shear stress experiments

Human umbilical vein endothelial cells (HUVECs) were plated onto a 6 well dish at 40000 cells/well. 24h later cells were transfected with TransIT-2020 (Mirus, MIR 5404) as previously described (Nawara et al., 2022) using 250 μl of DPBS (Sigma, D8537), 1 μg of CLCa-STAR plasmid, and 1 μl of RT TransIT-2020. Media was replaced after incubation with transfection mixture for 6 h at 37 °C and 5% CO_2_. 24h later expression of the plasmid was confirmed by epifluorescence and cells from all 6 wells were collected by trypsinization and spun down (150g for 5 min). The cell pellet was then resuspended in 200 μl of fresh media and 100 μl of cell suspension was plated per µ-Slide I Luer Glass Bottom (Ibidi, 80177). Before plating the cells, 200 μl of fresh media was added to the µ-Slides and allowed to equilibrate in the incubator. After cell attachment, 60 μl of fresh media was added to each of the µ-Slide reservoirs. 3-4 h later, slides were gently flushed with 1ml of media. 24h after plating, cells were again flushed with 1 ml of media. Then the flow system was assembled. Here we used a peristaltic pump (Fisherbrand, 13-876-2 or Kamoer Fluid Tech, DIP-B253) to circulate media. Briefly, tubing (Ibidi, 10831), and 10 ml syringe (Fisherbrand, 14-955-459) were preheated to 37°C. Then the tubing was connected to disinfected peristaltic pump silicone tubing. The plunger from a 10 ml syringe was removed and syringe was inserted into one of the µ-Slide inlets. The syringe serves as a media reservoir and air bubble trap. Then the tubing was connected to the second µ-Slide inlet and the end of the assembled was placed inside the syringe. This was followed by a gentle addition of 10 ml of media to the syringe. The top of the syringe was then parafilmed to mount the tubing and seal the system from contamination. Everything was placed into the incubator for at least 30 min to allow for equilibration. After that silicon tubing was mounted into the pump and flow was set to achieve shear stress of 10 dyn/cm^2^. The shear stress was then calculated according to manufacture protocol. Cells plated on the second µ-Slide were used as static control. 24h later cell alignment with the flow direction was confirmed. Then the µ-Slide was disconnected from the flow, followed by a media change to a media with biliverdin. 30 min later media containing biliverdin was washed and replaced with regular media and live cells experiments were performed as previously published under static conditions (Nawara et al., 2022).

### Image processing

Data were corrected and analyzed using Fiji (ImageJ, National Institutes of Health, Bethesda, MD), MATLAB version 2018b for CMEanalysis, and MATLAB 2020b for DrSTAR.

### Statistics and reproducibility

Data was acquired from three independent replicates and total of n_Stat_ = 21 cells and n_FSS_ = 18 cells. Figure legends contain description of statistical test used and p values. All results are presented as mean ± SEM unless otherwise noted. Statistical calculations were performed in GraphPad Prism (Version 9.0.0).

## Results

### Clathrin dynamics are increased in HUVECs cultured under FSS

To investigate CME in physiological vasculature conditions, we focused on the impact of biophysical stresses generated by blood flow on clathrin dynamics in ECs. HUVECs were cultured under static or fluidic shear stress (FSS; 10 dyn/cm2) conditions for 24 h. As expected, FSS caused overall morphological changes and the alignment of HUVECs with the direction of the flow (**Fig. 1A**). In HUVECs exposed to FSS we also observed reorganization of the actin cytoskeleton compared to control HUVECs, with alignment of actin fibers parallel to the direction of flow, similar to previous reports (Dean et al., 2023; Noria et al., 2004) (**Fig. 1B**). Next, we transfected HUVECs with clathrin light chain a (CLCa) tagged with the STAR probe (iRFP713-EGFP fusion). Surprisingly, live-cell TIRF microscopy revealed that HUVECs exposed to FSS have increased plasma membrane-associated clathrin dynamics (**Fig. 1C, D**).

**Fig. 1.**
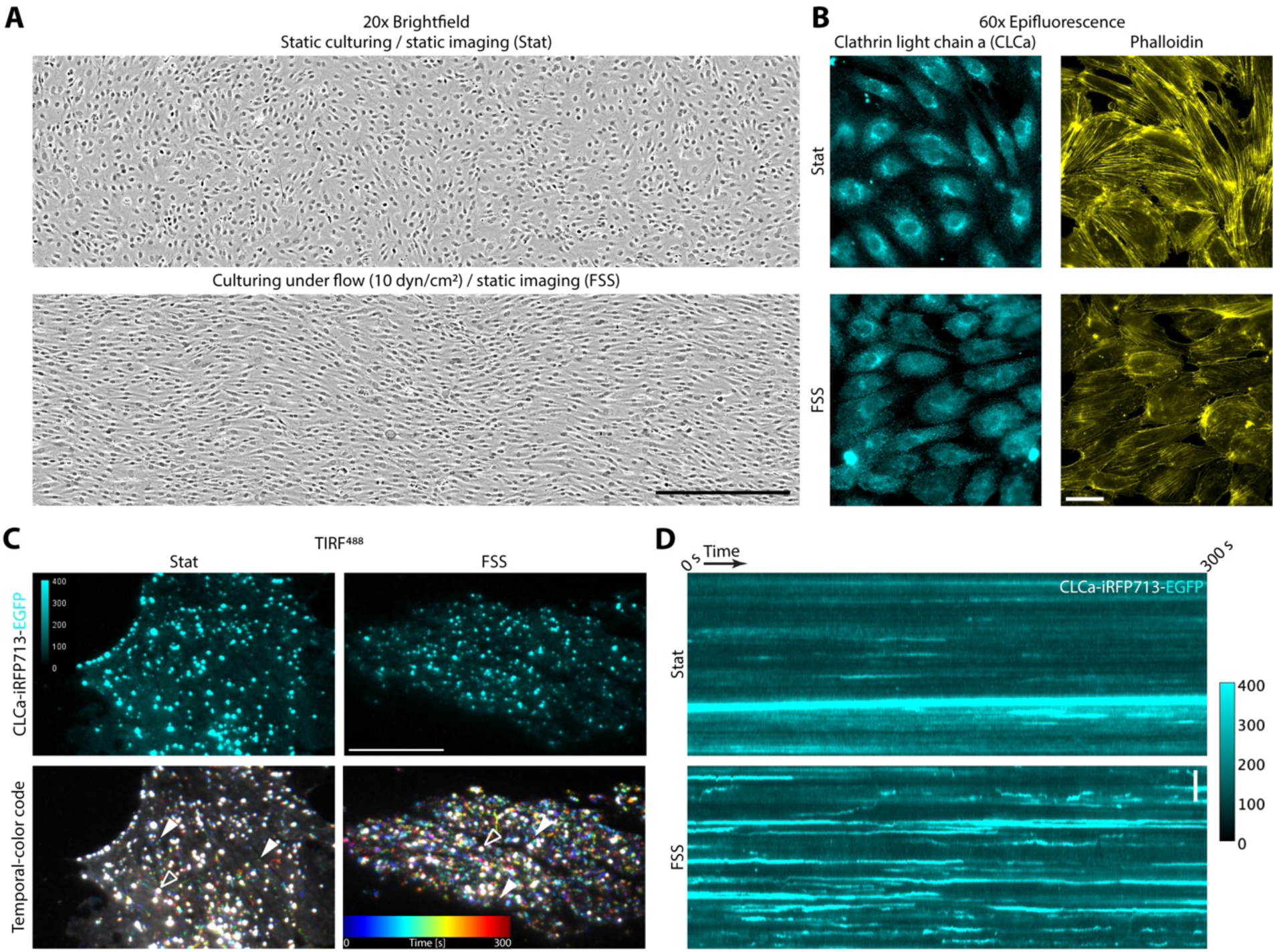
Clathrin dynamics are increased in HUVECs cultured under FSS. (**A**) Brightfield images of HUVECs grown under static (Stat - top) and FSS (10 dyn/cm2 - bottom) conditions for 24h, scale bar = 500 µm. (**B**) Clathrin light chain a and actin distribution in HUVECs grown under static or FSS conditions, scale bar = 40 µm. (**C**) Live-cell TIRF imaging shows increased clathrin dynamics in HUVECs grown under FSS compared to static (representative cells), scale bar = 20 µm. Temporal-color code projects a 5-minute-long movie onto a single image. Each frame is color-coded for time. *De novo* clathrin accumulations are multi-colored with the specific color based on the time of appearance and lifetime and (full carets) while persistent clathrin structures are white (empty carets). (**D**) Kymograph showing clathrin accumulation (intensity, [a.u.]) for HUVECs cultured under static or FSS conditions, scale bar = 5 µm.

### Both vesicle formation and flat clathrin accumulations are increased in HUVEC exposed to FSS

Conventional TIRF microscopy provides excellent time resolution for studying CME dynamics, however it is virtually impossible to separate productive clathrin accumulations from non-productive based on their fluorescent signal (Nawara and Mattheyses, 2023). To overcome this limitation we used STAR microscopy to parse out whether the observed increase in clathrin dynamics was caused by the surge in formation of productive clathrin-coated vesicles or nonproductive flat clathrin lattices (**Fig. 2A**) (Nawara et al., 2023; Nawara and Mattheyses, 2023; Nawara et al., 2022). STAR uses a ratiometric approach to recover axial Δz information, which requires the dual-tag (CLCa-iRFP713-EGFP). The bespoke MATLAB-based data processing pipeline DrSTAR was employed to process multiple cells and experimental replicates in a robust and high throughout manner (Nawara et al., 2023). Next CMEanalysis tracked clathrin events automatically based on their florescence signal allowing quantification of hundreds of clathrin events (Aguet et al., 2013). This was followed by classification of all diffraction-limited *de novo* clathrin accumulations as productive endocytosis (clathrin-coated vesicles - positive Δz) or as flat clathrin lattices (FCLs - no change in Δz; **Fig. 2B, C**). When productive clathrin events were divided into cohorts based on their total lifetime, we observed a similar initial rise of Δz for all cohorts in cells grown under static conditions (**Fig. 2B** – purple dashed rectangle). However, in cells exposed to FSS we observed a correlation between the lifetime of the clathrin accumulation and the rate of change in Δz (**Fig. 2C** – purple dashed rectangle). Longer-lived clathrin events formed vesicles at a slower rate in cells grown under the FSS. We had hypothesized that FSS may have a similar impact as mechanical stressors and cause disruption or inhibition of CME, which would result in an increase in flat clathrin lattices but not vesicle formation. Contrary to our hypothesis, we observed a 2.3-fold increase in CCV formation and a 1.9-fold increase in non-productive flat clathrin lattices in HUVECS grown under FSS compared to cells grown in static conditions (**Fig. 2D, E**). These data show that the increased clathrin dynamics in HUVECs grown under FSS represent both clathrin vesicle formation and flat clathrin lattices. This shows FSS is a mechanical stimulus of CME in HUVECS, contrary to most mechanical stresses evaluated in other cellular contexts.

**Fig. 2.**
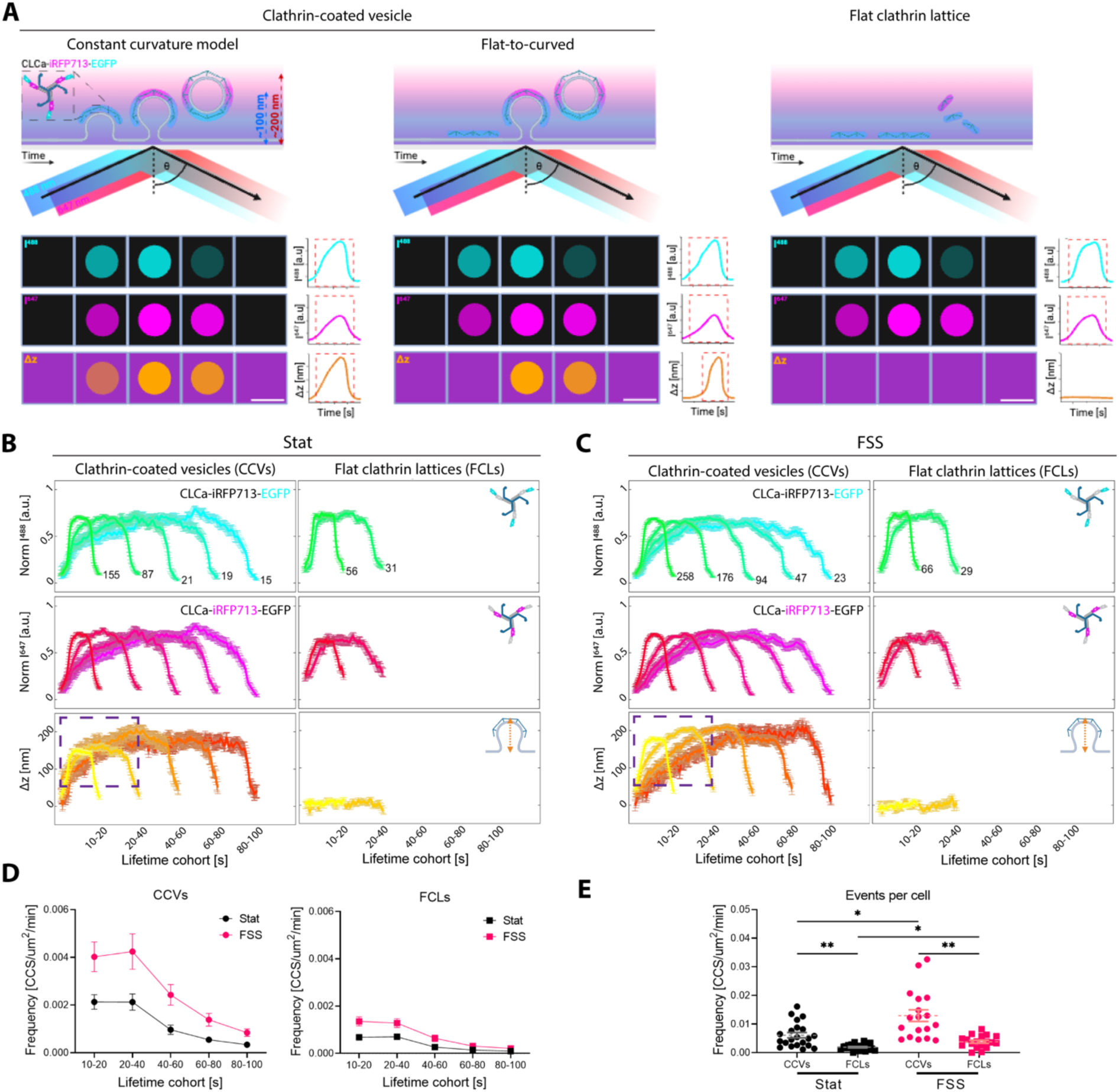
More productive and flat clathrin accumulations are observed in HUVEC exposed to FSS. (**A**) STAR microscopy can identify productive clathrin-coated vesicles and unproductive flat clathrin lattices. CLCa-iRFP713-EGFP (STAR probe) imaged by STAR during: clathrin-coated vesicle formation following the constant curvature (left) or flat-to-curved models (middle) and flat clathrin lattice assembly (right). Each event is depicted as a schematic of the images that will be acquired (top panels) and a schematic of a single endocytic puncta as imaged by the camera (panels labeled I^488^ (cyan) – intensity at 488 nm and I^647^ (magenta) – intensity at 647 nm) as well as the calculated in a pixel-by-pixel manner Δz from the I^647^/I^488^ ratio. On the right from the camera view are “measured” intensity and Δz traces for the example clathrin accumulations. Those single traces are then pulled into lifetime cohorts as in **B** and **C**. Evanescent wave penetration depths (dashed arrows) correspond to 488 (blue) and 647 (red) nm lasers where ϴ is the incidence angle. Scale bar = 100nm. Formation of CCVs or flat clathrin lattices (FCLs) in HUVECs grown under static (Stat - **B**) and FSS (10 dyn/cm2 - **C**) conditions for 24h was determined using simultaneous two-wavelength axial ratiometry (STAR) microscopy. N of representative clathrin accumulations for each lifetime cohort are indicated (mean ± SEM; EGFP - cyan, iRFP713 - magenta, Δz which indicates vesicle formation - orange). The dashed square highlights the first 0 - 40 s of vesicles formation. (**D**) Histogram of lifetime distribution of clathrin-coated vesicles (CCVs) and flat clathrin lattices (FCLs) per µm^2^, per minute (mean ± SEM). (**E**) Cumulative frequency of CCVs and FCLs per µm2, per minute (mean ± SEM, * - p<0.03, ** - p<0.002, Brown-Forsythe and Welch ANOVA tests followed by Dunnett’s T3 multiple comparisons test). Data was acquired from three independent replicates and total of n_Stat_ = 21 cells and n_FSS_ = 18 cells.

### Initiation of clathrin-coated vesicle formation is delayed after exposure to FSS

We next asked whether the mechanism of CCV formation is altered in response to FSS. To do so, we quantified the initiation of CCV formation relative to clathrin arrival, reported as the time difference in when Δz and CLCa are detected (Nawara et al., 2023; Nawara et al., 2022). Multiple productive routes of clathrin-coated vesicle formation have been observed in living cells, which have been classified into three main models (Chen and Schmid, 2020; Nawara et al., 2022; Scott et al., 2018). The nucleation model assumes that plasma membrane curvature occurs before clathrin recruitment. This can be mediated by curvature sensing/inducing proteins, receptor clustering etc. (Bhave et al., 2020). In the constant curvature model, there is simultaneous clathrin recruitment and vesicle formation (**Fig. 2A, left**). Finally, the flat-to-curved transition model is based on preassemblies of flat clathrin lattices that then dynamically bend to form a vesicle (**Fig. 2A, right**) (Nawara et al., 2022).

When all productive endocytic events were analyzed, it revealed a significant delay in the initiation of vesicle formation relative to when clathrin was first observed in HUVECs exposed to FSS (**Fig. 3**). We hypothesized that preassembling stable flat clathrin lattice before vesicle formation might be preferential under stress. This could prevent clathrin-coated structure or plasma membrane rupture during stress. Therefore, we next asked if this delay was dependent on event lifetime. We found that in cells grown under FSS, only clathrin events longer than 50 s favor the pre-assembly of flat clathrin lattices and the flat-to-curved mode of CCV formation (**Fig. 4** **A, B**). This suggests that preassembling a flat clathrin platform before vesicle initiation is a preferred mechanism of CCV formation under conditions in the vasculature.

**Fig. 3.**
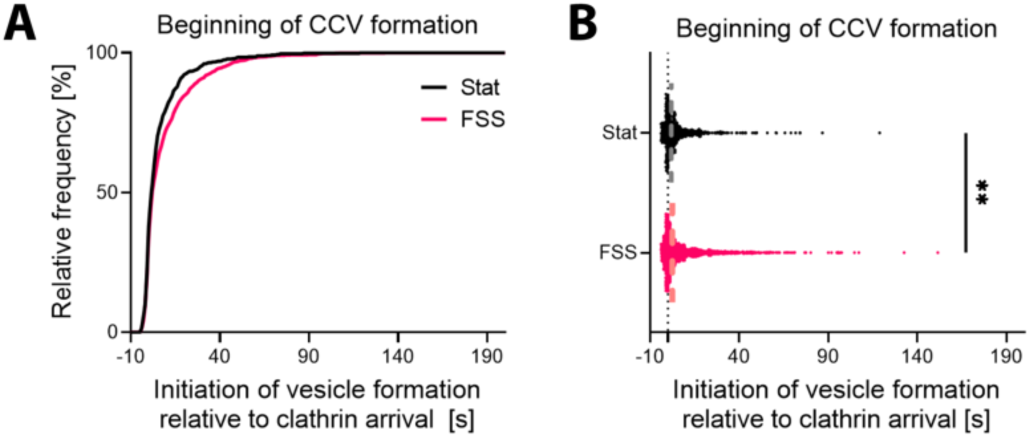
Initiation of clathrin-coated vesicle formation is delayed after exposure to FSS. (**A**) Cumulative distribution of the initiation of vesicle formation (Δz) relative to the detection of CLCa-STAR, reported as reported as Δz_Beg_ - CLCa_Beg_. (**B**) Distribution of the initiation of vesicle formation (dashed line indicates median, ** - p=0.09, Kolmogorov– Smirnov test). Data was acquired from three independent replicates and total of n_Stat_ = 21 cells and n_FSS_ = 18 cells.

**Fig. 4.**
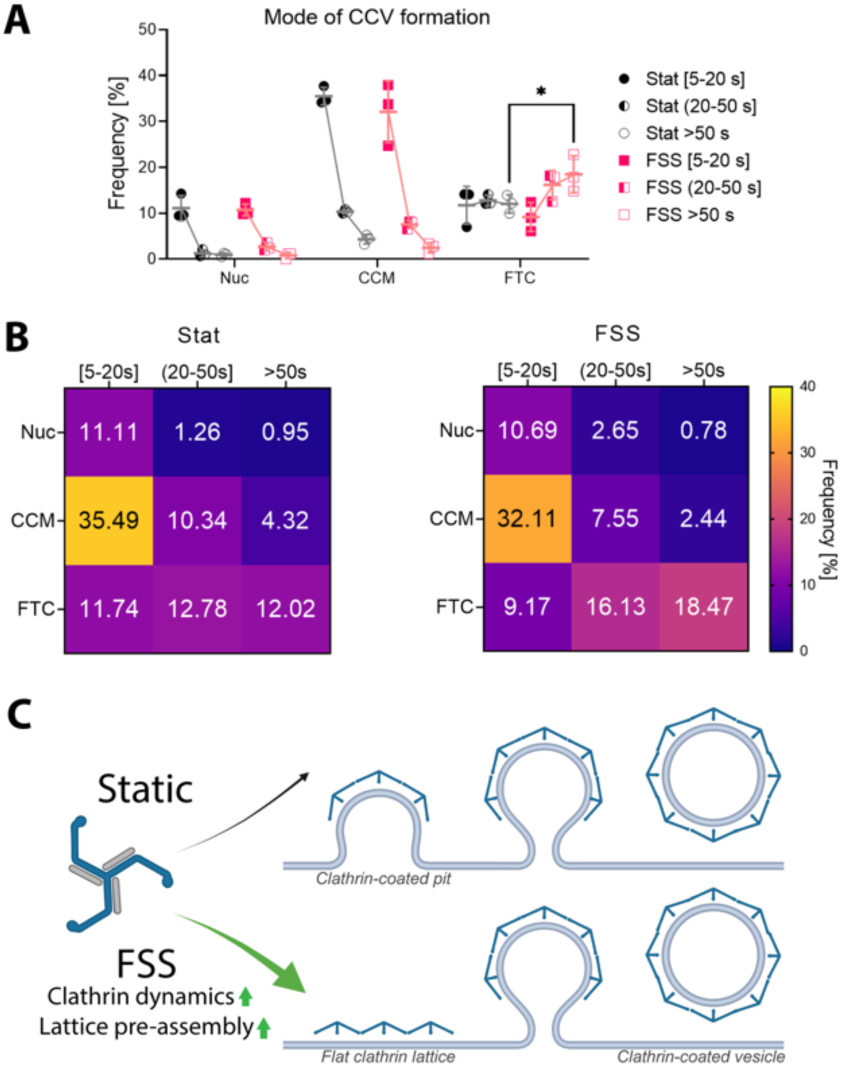
Mode of clathrin-coated vesicle formation is changed following FSS. (**A**) Distribution of events across three models of vesicle formation and three lifetime cohorts in HUVECs grown under static (Stat, black) and FSS (10 dyn/cm^2^; magenta), Nuc = Nucleation, CCM = Constant Curvature Model, FTC = Flat-to-curved transition. (mean ± SD, * - p = 0.048 by two-way ANOVA followed by Tukey’s multiple comparisons test) (**B**) Means from (**A**) reported as a frequency heat map for static (left) and FSS (right). Data was acquired from three independent replicates and total of n_Stat_ = 21 cells and n_FSS_ = 18 cells. (**C**) FSS upregulates clathrin dynamics and promotes FTC vesicle formation in HUVECs.

## Discussion

We have shown that FSS increases clathrin dynamics and modulates the mechanism of CCV formation in ECs (**Fig. 4C**). It is possible that both changes work to preserve robust endocytosis in the vasculature by countering the inhibitory effects of cyclic stretch, plasma membrane tension and other mechanical stresses (Dessalles et al., 2021; Joseph and Liu, 2020). One explanation for the increase in membrane associated clathrin dynamics is an upregulation of endocytic accessory proteins. Interestingly, a more peripheral clathrin signal was observed in HUVECs exposed to FSS when imaged with epifluorescence (**Fig. 1B**). This leads us to propose that alternatively clathrin itself could be redirected to the plasma membrane as a direct result of FSS (Charwat et al., 2018). This localization could be achieved, for example, by activation of fast-loop recycling of endosomes mediated by Rab4 (Kofler et al., 2018). Our data shows that HUVECs exposed to FSS have slower vesicle formation dynamics (**Fig. 2A, B** and **Fig. 3**). Moreover, longer events (>50 s) prefer to preassemble a stable flat clathrin platform before initiation of vesicle formation (**Fig. 4A, B**). Both of those phenomena may be protective against plasma membrane injury during cargo internalization. We hypothesize that different levels of FSS as well as different flow patterns may require ECs to adapt accordingly. This work opens the door to many interesting questions about the nature of such adaptation. Are endocytic accessory proteins upregulated or redistributed? Are there any markers that highlight the productive or unproductive sites? Is the increase of clathrin dynamics related to reorganization of cytoskeleton?

This study highlights the importance of considering the physiological local vascular environment in the context of internalization of growth factors, membrane proteins, therapeutics, or pathogens. Studies in non-ECs or ECs not cultured under FSS may not properly recapitulate endocytosis rates or clathrin dynamics and could lead to incorrect conclusions. Further research into the biophysical framework of CME in ECs under physiological and stress conditions will provide critical insight required for targeted drug delivery to promote cardio-vascular health and treat disease (Jones and Shusta, 2007). As more tools are developed to allow quantification of endocytosis in primary cells in physiological environments it will prove key to revisit principles derived from continuous cell culture experiments (Chan et al., 2022; Schöneberg et al., 2018; Sochacki et al., 2021).

## Author contributions

Conceptualization: T.N.; methodology: T.N. and A.M.; validation: T.N.; formal analysis: T.N.; investigation: T.N.; resources: T.N., E.S., and A.M.; data curation: T.N.; writing— original draft: T.N.; writing—reviewing and editing: T.N., E.S. and A.M.; visualization: T.N.; supervision: E.S. and A.M.; funding acquisition: T.N. and A.M.

## Acknowledgments

The authors support diversity and inclusiveness in science and believe strongly in equality of human rights. The authors would like to thank Andrew Kowalczyk (Penn State) and Aleksandra Michrowska (The Francis Crick Institute), as well as members of the Mattheyses and Sztul laboratories for their support and helpful discussion. We thank UAB IT Research Computing. This work was supported by American Heart Association 906086 to T.N., NIH/NIGMS R01GM131099 to A.L.M. and NSF CAREER 1832100 to A.L.M.

